# SoyDNGP: A Web-Accessible Deep Learning Framework for Genomic Prediction in Soybean Breeding

**DOI:** 10.1101/2023.06.15.545107

**Authors:** Pengfei Gao, Haonan Zhao, Zheng Luo, Yifan Lin, Yaling Li, Fanjiang Kong, Chao Fang, Xutong Wang

## Abstract

Soybean is a globally significant crop, playing a vital role in human nutrition and agriculture. Its complex genetic structure and wide trait variation, however, pose challenges for breeders and researchers aiming to optimize its yield and quality. Addressing this biological complexity requires innovative and accurate tools for trait prediction. In response to this challenge, we have developed SoyDNGP, a Convolutional Neural Networks (CNN)-based model that offers significant advancements in the field of soybean trait prediction. Compared to existing methods, such as DeepGS and DNNGP, SoyDNGP boasts a distinct advantage due to its lower parameter volume and superior predictive accuracy. Through rigorous performance comparison, including prediction accuracy and model complexity, SoyDNGP consistently outperformed its counterparts. Furthermore, it effectively predicted complex traits with remarkable precision, demonstrating robust performance across different sample sizes and trait complexities. We also tested the versatility of SoyDNGP across multiple crop species, including Cotton, Maize, Rice, and Tomato. Our results showed its consistent and comparable performance, emphasizing SoyDNGP’s potential as a versatile tool for genomic prediction across a broad range of crops. To enhance its accessibility to users without extensive programming experience, we have designed a user-friendly web server, available at http://xtlab.hzau.edu.cn/SoyDNGP. The server provides two primary features: ‘Trait Lookup’, offering users the ability to access pre-existing trait predictions for over 500 soybean accessions, and ‘Trait Prediction’, allowing for the upload of VCF files for trait estimation. By providing a high-performing, accessible tool for trait prediction and genomic analysis, SoyDNGP opens up new possibilities in the quest for efficient and optimized soybean breeding.

## Introduction

Food insecurity is a growing issue, heightened by the increasing world population and the challenges of climate change (UNICEF, 2021). Improving crop yield, especially in crucial crops like soybean, is critical. Traditional breeding methods, while effective, can be slow and struggle to keep pace with the demands. For instance, they fall short of achieving the annual yield improvement rate of 2.4% needed to double global soybean production by 2050 (Ray *et al*., 2012; Yoosefzadeh-Najafabadi *et al*., 2022). To speed up the process, breeders have turned to genomic tools, including genomic selection (GS) (Decker, 2015).

GS is an advanced technique that can make breeding faster and more efficient (Bhat *et al*., 2016). It uses genomic prediction models, along with many genetic markers across the genome, to predict how a trait will perform (Meuwissen *et al*., 2001; Hayes & Goddard, 2007; Sandhu *et al*., 2021; Wang *et al*., 2023). It has been used successfully in both animal and plant breeding, especially in improving traits like crop yield, breeding value, and disease resistance (Meuwissen *et al*., 2001; Desta & Ortiz, 2014; Bhat *et al*., 2016; Poland & Rutkoski, 2016). However, using GS effectively relies on many factors, like the size of the training population, heritability of traits, marker density, and the prediction model used (Xu & Crouch, 2008). Traditional models, such as linear regression models (GBLUP, rrBLUP, and Bayesian methods), often struggle with capturing complex non-additive effects (Meuwissen *et al*., 2001; VanRaden, 2008; De Los Campos *et al*., 2009; Endelman, 2011). That’s where deep learning methods, such as DeepGS and DNNGP, can play a role (Ma *et al*., 2017; Wang *et al*., 2023). They use multiple hidden layers to capture complex, non-linear relationships in the data. However, these techniques require large datasets for accurate predictions, which can be a challenge in some cases.

Soybean [*Glycine max* (L). Merr.] is a globally significant crop, providing a rich source of protein and oil for human and animal consumption (Hartman *et al*., 2011). In soybean breeding, genomic prediction has already shown its potential (Stewart-Brown *et al*., 2019; Ravelombola *et al*., 2021; Yoosefzadeh-Najafabadi *et al*., 2022). It is crucial in improving soybean varieties to meet the increasing demand and to tackle the challenges posed by climate change, pests, and diseases. But, there are still hurdles to overcome, like capturing the full range of genetic diversity in soybean and refining genomic prediction methods.

In this study, we aim to look more closely at how deep learning methods can be used for genomic prediction in soybean breeding. We used a rich source of soybean genomic data, like the genotypes of thousands of soybean samples from the USDA Soybean Germplasm Collection and hundreds of phenotypes from the GRIN-global web server (Postman *et al*., 2009; Song *et al*., 2015). We remodeled a deeper neural network framework for our method, called SoyDNGP. We then compared the accuracy of prediction in SoyDNGP with the deepGS approach for several important traits. We also test SoyDNGP with other species to check if it’s compatible beyond soybeans. Finally, to make SoyDNGP accessible to a wide audience, we’re building a user-friendly web server that provides a comprehensive approach to refining genomic predictions and doesn’t require any programming knowledge. Our study hopes to shed light on how deep learning methods can be applied in soybean breeding and contribute to improving genomic selection strategies.

## Materials and methods

### Datasets used for genomic prediction of soybean

The data used for training and predicting with the SoyDNGP model were procured from two comprehensive online databases: SoyBase and GRIN-Global(Postman *et al*., 2009; Grant *et al*., 2010). The genotype information of an impressive collection of 20,087 soybean accessions, including 42,509 high-confidence SNPs (Single Nucleotide Polymorphisms) based on the SoySNP50K iSelect BeadChip, was sourced from SoyBase. The pre-built Variant Call Format (VCF) file, corresponding to version 2 of Williams 82 reference sequences, was utilized. In an attempt to enhance the compatibility of our model with SNP datasets derived from methods other than the 50K SNP chip, an intersection operation was performed with SNP loci generated from resequencing data. The Beagle 5.4 program (version 22Jul22.46e) was used to phase the SNPs and fill in the missing data (Ayres *et al*., 2012). This process resulted in a curated set of 32,032 SNP loci, which were used for model training. However, to maintain a uniform set of annual species and minimize the component of mixed accessions, a selection process was implemented, reducing the number of soybean accessions used for model construction to 13,784. These chosen accessions, part of the USDA Germplasm Collection, are representative of a broad spectrum of landraces and elite cultivars from around the globe.

Phenotypic data for each of these selected soybean accessions were obtained from the GRIN-Global database (https://npgsweb.ars-grin.gov/gringlobal/search). Despite an initial collection of 23 agronomic traits, our focus was narrowed down to ten key traits. This included six quantitative traits such as protein content, oil content, hundred-seed weight (SdWgt), flowering date (R1), the maturity date (R8), yield, and plant height (Hgt). In addition, four qualitative traits were also considered, which encompassed stem termination (ST), flower color (FC), pubescence density (PDENS), and pod color (POD).

### SoyDNGP model structure

In stark contrast to the traditional DNNGP’s three-layer wide convolution architecture used for genome-wide big data analysis, our SoyDNGP utilizes a deep and slim network structure (Wang *et al*., 2023). This structure is inspired by the concept of segmentation drawn from the VGG deep learning network (Simonyan & Zisserman, 2014). Specifically, SoyDNGP is built around ‘convolutional blocks’, each incorporating a convolutional layer, normalization layer, and an activation layer (ReLU).

Every feature extraction unit in the network is comprised of one or two of these convolutional blocks, resulting in an effective block structure for feature extraction. At the end of the convolutional sequence, we’ve included a fully connected layer to enhance the expression capabilities of the network. With the network’s increased depth, we’ve also added a dropout layer (dropout = 0.3) after each convolution to mitigate overfitting and a normalization layer to enhance the model’s ability to generalize. Overall, the network architecture integrates 12 convolutional layers and a single fully connected layer, designed to handle an input tensor of dimensions (206 × 206 × 3).

The first convolutional module operates using a 3 × 3 convolution kernel with a stride of 1, which effectively upscales features and expands the feature map from 3 channels to 32. The subsequent convolutional block deploys a 4 × 4 convolution kernel with a stride of 2, increasing the dimensions of the feature map while simultaneously reducing the size of each dimension’s feature map. In the network structure that follows, each feature extraction block consists of two convolutional layers.

In each feature extraction block, the first convolutional layer adjusts the convolution kernel size and sampling stride based on the dimensions of the feature map. This guarantees a complete traversal of the feature map while enabling feature map scaling and dimensionality increase with the smallest feasible convolution kernel. The second convolutional layer uses a 3 × 3 convolution kernel to reprocess the feature map from the preceding layer, reinforcing feature extraction. This process is iterated until the feature map’s channel count escalates to 1024, with dimensions reducing to 7 × 7. Subsequently, the feature map is flattened into a one-dimensional vector, and forwarded to the fully connected layer for final classification and regression processing. Given the extensive information density of the SNP variation-based feature matrix, we have chosen to move away from the simplistic zero-padding approach during convolution padding. Instead, we apply a symmetrical filling technique that leverages matrix elements at the outermost layer, using the matrix edge as the axis of symmetry. This significantly bolsters the feature extraction capability from the matrix.

To circumvent the potential issue of overfitting in the model training process induced by the depth of the network, weight decay was applied to the Adam optimizer. This included a decay rate of 1e-5 for regression and 0.01 for classification. For qualitative traits, the model was trained using the commonly applied cross-entropy loss function. Conversely, for regression tasks pertaining to quantitative traits such as protein content and yield, SoyDNGP utilized the Smooth L1 Loss function (β = 0.1) as its loss function:

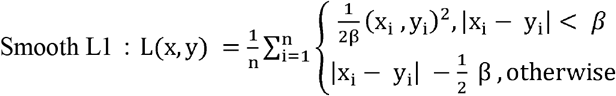

This particular loss function provides a constant gradient when the loss is significant, thereby mitigating the potential disruption of training parameters due to substantial gradients. Conversely, when the loss is minimal, the gradient dynamically reduces, addressing the challenge of convergence often seen with L1 loss. Compared to traditional L1 and L2 loss functions, the Smooth L1 Loss function offers accelerated convergence speed, improved robustness to outliers, and enhanced gradient smoothness. For each trait under consideration, we conducted 150 epochs on GeForce RTX 3090 or RTX A6000, selectively preserving the epoch that demonstrated optimal performance on the test set as the final model weights.

Lastly, it’s noteworthy that we’ve incorporated a Coordinate Attention (CA) mechanism module after the first and final convolutional layers (Hou *et al*., 2021). This strategy amplifies attention to the positional information in the feature matrix and between channels, thereby enhancing spatial information extraction. SoyDNGP’s model structure is designed and implemented using PyTorch (version 2.0.1), a widely recognized open-source machine learning library (Imambi *et al*., 2021).

### Remodeling DeepGS in Python

To facilitate a fair comparison between model architectures, and acknowledging the limited feature representation ability of the original DeepGS model, we opted to enhance it while maintaining its overall structure (Ma *et al*., 2017). This structure comprises a combination of a convolution layer, a ReLU activation function, a max pooling layer, and a dropout layer, all linked to two fully connected layers.

In the original DeepGS model, we substituted the broad 1 × 18 convolution kernel with a more compact 3 × 3 kernel. Additionally, we increased the count of both convolution and pooling layers in the model to six, yielding a total of twelve layers. This modification ensured the channel count of the final feature map aligned with that of SoyDNGP. After adjusting the model structure, we preserved all other conditions identical to those in the SoyDNGP model for the training phase. This approach enabled us to draw an equitable comparison between the two model architectures. Moreover, it underscored the superiority of a slender and deep convolutional network in the realm of feature extraction and representation capabilities.

### Model construction of traditional machinal learning algorithms

In order to gauge the effectiveness of our proposed SoyDNGP model, we conducted parallel evaluations using nine conventional machine learning algorithms on identical datasets. These traditional models encompassed: K-Nearest Neighbors (KNN), Decision Tree (DT), Random Forest (RF), Multilayer Perceptron (MLP), Adaptive Boosting (Adaboost), Gaussian Naive Bayes (GNB), and Support Vector Classification (SVC) with different kernels—Linear, Radial Basis Function (RBF), and Sigmoid (Hsu *et al*., 2003; Myles *et al*., 2004; Peterson, 2009; Biau & Scornet, 2016; Ramchoun *et al*., 2016; Ontivero-Ortega *et al*., 2017; Feng *et al*., 2020). Each trait was subjected to training using these nine algorithms, facilitating a comparative analysis of their performance and robustness against SoyDNGP on the same dataset. The hyperparameter configurations for these models were as follows: In KNN, we assigned the number of neighbors (n_neighbors) as 3. For DT and RF, we confined the maximum depth of the trees (max_depth) to 5, while for RF, we also defined the number of trees in the forest (n_estimators) as 10, and the number of features considered for the optimal split (max_features) as 1. For MLP, we stipulated the L2 penalty (regularization term) parameter (alpha) as 1. The remaining models utilized their default parameters as defined in their respective libraries.

We implemented a 10-fold cross-validation scheme (n_splits=10) for a more rigorous evaluation of the models, ensuring diverse splits for each run (random_state=None), and random shuffling of the data prior to fold creation (shuffle=True). This was done to preclude the possibility of any class’s overrepresentation in any given fold, which might skew the model’s performance. Our assessment metrics consisted of precision, recall, and F1-score for each class of traits. Additionally, we calculated the mean and standard deviation of accuracy across the folds, offering an encompassing view of the model’s performance. To ascertain their generalizability, we evaluated the models based on their accuracy on both training and test datasets.

### Data processing of resequencing of soybean database

To assess the performance of SoyDNGP in different soybean populations, we obtained resequencing data from two public datasets available on NCBI and GSA. The following steps were taken to process all sequencing reads: Initially, Cutadapt (version 3.5.0) was employed to excise potential adaptors and discard low-quality reads(Martin, 2011). The clean reads were then aligned to the version 2 of the Williams 82 reference genome sequences (https://phytozome-next.jgi.doe.gov/) utilizing BWA (version 0.7.17-r1188) (Li, 2013). Next, PCR duplicates and reads that mapped to multiple locations were eliminated using Samtools (version 1.15.1-41-gc7acf84) (Li, 2013; Danecek *et al*., 2021). The GATK pipeline was subsequently deployed to produce reliable SNPs for variation and evolutionary analysis. SNPs were preserved if they were bi-allelic and had a MAF greater than 0.05 (McKenna *et al*., 2010). To maintain SNP loci that overlapped with the SoyDNGP training data, a total of 32,624 SNPs were chosen using VCFTools (version 0.1.16) (Danecek *et al*., 2011). Lastly, the Beagle 5.4 program was used to phase the SNPs and fill the missing data.

The VCF files generated were employed to predict the phenotypes using pre-built SoyDNGP models. We focused on a population comprising 559 soybean accessions for which three phenotypes, namely flowering time, hundreds of seeds weight, and plant height, had been previously measured. These phenotypes were recorded in 2018 for plants cultivated in Zhengzhou, Henan Province, China (latitude 34.7N, longitude 113.6E). To quantify the accuracy of the phenotype predictions, we computed the Pearson correlation coefficient (*r* value) between the observed phenotypes and the predicted values.

### Web server implementation

Our web server is established on a Webflow template (https://webflow.com/), which is enhanced with the Bootstrap5 framework (https://getbootstrap.com/) and operated via the Flask web framework (https://flask.palletsprojects.com/) (Hammer *et al*., 2009). To facilitate additional functions, we’ve chosen several specific tools. Redis (https://redis.io/) functions as the custodian of progress data and prediction results, while MongoDB (https://www.mongodb.com/) is employed to store the data available for users (Banker *et al*., 2016; Gade *et al*., 2018). Gunicorn (https://gunicorn.org/), a Python WSGI HTTP server, manages server operations, and Nginx (https://nginx.org/en/) is used for request forwarding from port 80 to the Gunicorn service, as well as for load balancing (Reese, 2008). The entire project is hosted on a Linux-based system equipped with an i7-13700KF processor and an RTX 3060Ti graphics card. The components of SoyDNGP’s Web Server were illustrated in Figure S1.

## Results

### SoyDNGP exhibits impressive capabilities in soybean genomic prediction

SoyDNGP employs a three-dimensional layer input derived from standard VCF files. Given that there are three types of genotypes - reference base (0/0), homozygous variation (1/1), and heterozygous variation (0/1) - and in order to account for potential missing data (./.), SoyDNGP uses a phased process to fill in any gaps during training (Figure 1A). The input layer denotes these genotypes as follows: [1,0,0] for “0/0”, [0,1,0] for “0/1”, [0,0,1] for “1/1”, and [0,0,0] for “./.” genotypes (Figure 1A). This method allows the SoyDNGP structure to consider both the type of genotype and its spatial relationship. Two distinct structures are deployed for classification (qualitative traits) and regression (quantitative traits) tasks (Figure 1A).

**Figure 1.**
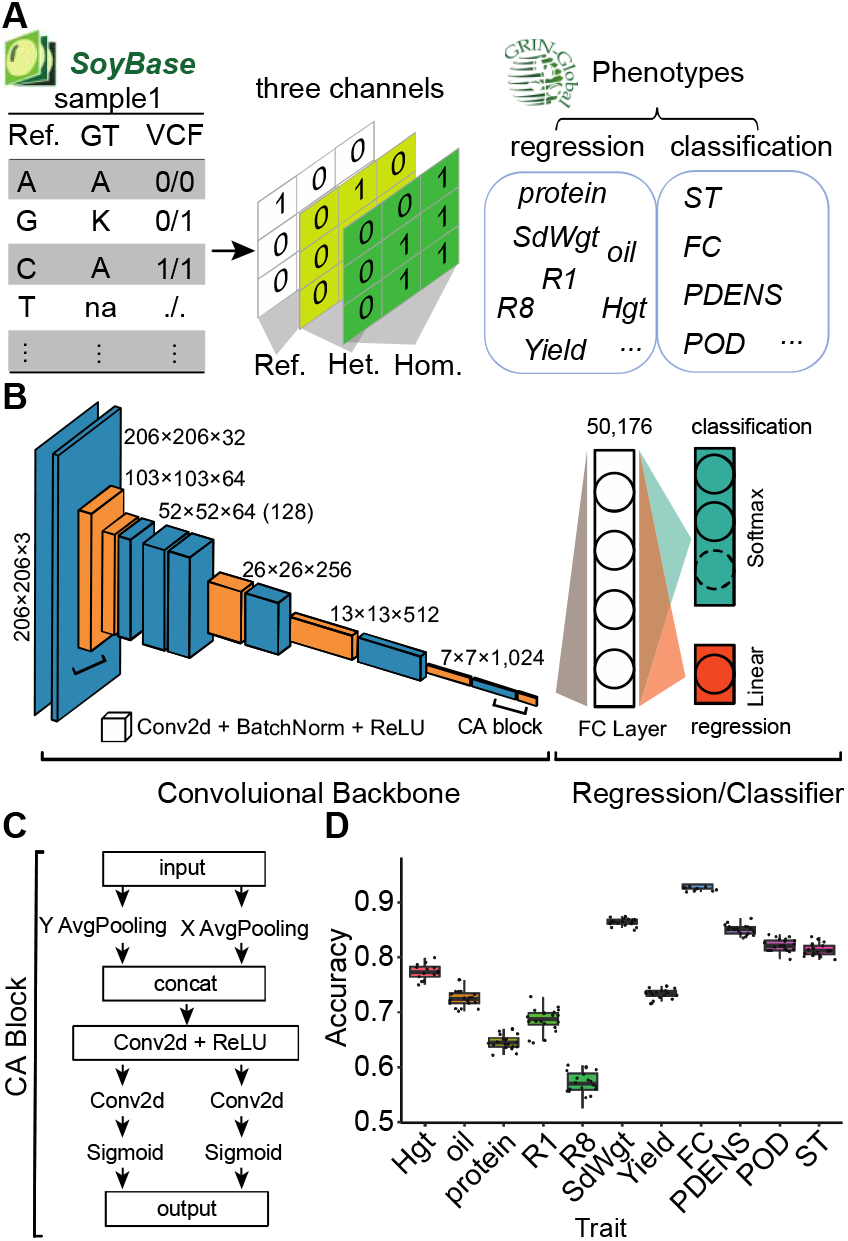
Overview of SoyDNGP’s Features. (A) The transformation process of genotype and phenotype data as input for SoyDNGP. (B) Depiction of the SoyDNGP module structure for classification and regression tasks. (C) Detailed illustration of the Coordinate Attention (CA) Block. (D) SoyDNGP’s predictive accuracy for eleven key agronomic traits.

SoyDNGP implements a Convolutional Neural Network (CNN) architecture that is characterized by twelve convolutional layers and one fully connected layer (Figure 1B). This structure employs a compact convolution kernel and a deep network design to efficiently perform feature extraction from the feature maps. During the training phase, the Adam optimizer (Adaptive Moment Estimation), which incorporates principles of Momentum and Adaptive Learning Rate methods, is utilized for updating the weights of the model. This optimizer strategy allows for efficient evasion from saddle points and accelerates the model’s convergence to optimal fitting. Coordinate Attention (CA) module is integrated following the initial and concluding convolutional layers. This technique amplifies focus on the spatial information within the feature matrix and inter-channel correlations, thereby boosting the extraction of spatial details (Figure 1C).

To ascertain the optimal sample size for model training, we trained the model using varying numbers of samples and monitored the predictive performance. The samples were divided into groups of 2k, 5k, 8k, and 10k for training, each paired with test sets of 11,784, 8,784, 5,784, and 3,784 samples respectively, over a span of 150 epochs. Our findings indicated that a sample size of 2k yields lower performance in terms of accuracy and other metrics, while no significant differences were observed among larger sample sizes (Figure S2 & S3). Ultimately, we found a sample size of 5k to be the most suitable for model construction.

We then conducted individual predictions to test accuracy. The results revealed that the prediction accuracy for regression tasks ranged from 0.56 in R8 to 0.87 in SdWgt, while for classification tasks it ranged from 0.82 in ST to 0.96 in FC (Figure 1D). These results are represented by the absolute errors between normalized observed and predicted phenotypic values as depicted in Figure S4. Through extensive testing, the model consistently delivers impressive prediction accuracy in both regression and classification tasks.

### Prediction accuracy of SoyDNGP and comparison with other algorithms

In order to evaluate the performance of SoyDNGP in genomic prediction relative to other machine learning (ML) methodologies, we utilized an identical dataset for training SoyDNGP models, which was also applied to other procedures. Although conventional machine learning techniques are not optimized for regression tasks, we discovered that several were capable of performing classification tasks with high accuracy. For instance, the Decision Tree model yielded prediction accuracies of 0.97 and 0.85 for FC and POD, respectively (Figure 2A). With the SVMRBF model, the accuracy of ST and PDENS reached 0.82 and 0.84 respectively. Among the nine ML methods tested, SoyDNGP exhibited balanced performance across all classification traits, with accuracies ranging from 0.82 (ST) to 0.94 (FC) (Figure 2A).

**Figure 2.**
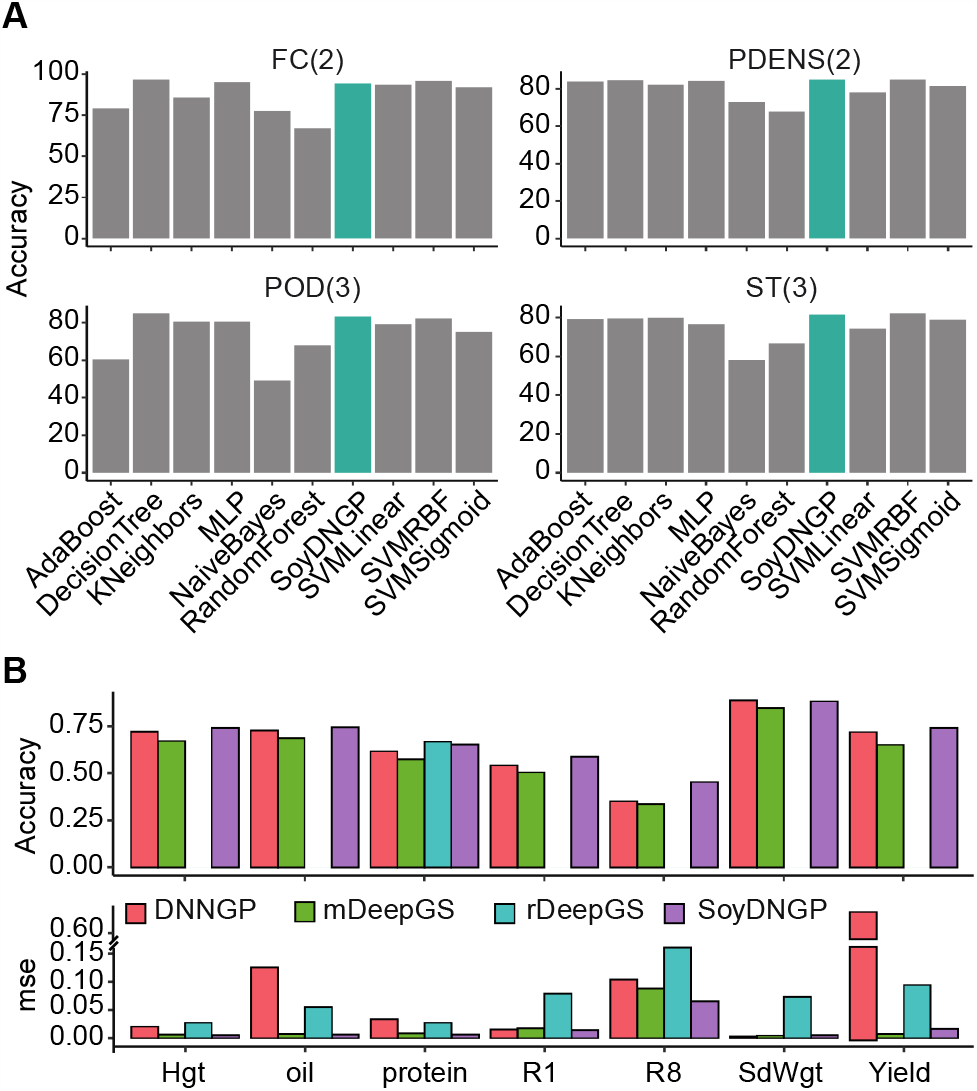
Comparative Analysis of Predictive Performance Between SoyDNGP and Other Approaches. (A) The predictive accuracy of SoyDNGP in comparison with traditional machine learning methods for classification tasks such as Flower Color (FC), Pubescence Density (PDENS), Pod Color (POD), and Stem Termination (ST). The numbers in parentheses on the plot denote the number of classification categories. (B) The predictive accuracy of SoyDNGP in comparison with other deep learning-based methods for regression tasks. Hgt, Oil, Protein, R1, R8, SdWgt, and Yield represent Plant Height, Oil Content, Protein Content, Flowering Time, Maturity Time, Hundred Seed Weight, and Yield, respectively. Accuracy is quantified by the correlation coefficient (r). ‘mse’ denotes Mean Standard Error, reflecting the absolute errors between the normalized observed and predicted phenotypic values.

To assess the performance of SoyDNGP in comparison with other CNN-based deep learning models, such as deepGS and DNNGP, we recreated their model architectures according to the details provided in the original research literature using Python. Regrettably, the original version of deepGS (rDeepGS) performed subpar and was unsuitable for regression tasks, despite its comparable performance in classification tasks with other methods (Figure 2B, Table 1). To confirm the efficiency of the deepGS structure, we re-engineered it into a modified version (mDeepGS). Training these models with the same dataset as used for SoyDNGP revealed that regardless of the trait or the amount of training samples utilized in regression tasks, SoyDNGP consistently outperformed both mDeepGS and DNNGP (Figure 2B). The correlation coefficient (r) for DNNGP deviated by approximately 5% from that of SoyDNGP (Figure 2B, upper panel). Moreover, the discrepancy between predicted and actual values (mse) was nearly ten times larger than with SoyDNGP (Figure 2B, lower panel). This suggests that DNNGP only has the capacity to predict trends and qualitatively describe them, but lacks precision in quantification. Owing to its shallow structure, mDeepGS was unable to effectively manage the complexity of the regression task, thus failing to fit accurately. Our observations revealed that the three deep learning models - DeepGS, DNNGP, and SoyDNGP - showed comparable performances for qualitative trait classification tasks. However, significant differences were observed in their performances on regression tasks. DeepGS, much like traditional machine learning models, failed to effectively fit the regression tasks. This could be primarily attributed to the less complex nature of classification tasks that can be effectively tackled with machine learning techniques, resulting in minimal differences in model performance across these tasks. Additionally, when compared with the three methods, the total parameter count in SoyDNGP is significantly lower, indicative of the model’s comparative simplicity (Table S1). This evidence indicates that the SoyDNGP model structure holds a significant advantage in genomic prediction compared to other methods.

**Table 1.** The predictive accuracy of SoyDNGP in comparison with other deep learning-based methods for classfication tasks.

| trait | methods | accuracy | precision | recall | f1 score |
| --- | --- | --- | --- | --- | --- |
| FC<sup>*</sup> | rDeepGS | 0.95 | 0.94 | 0.94 | 0.94 |
|  | mDeepGS | 0.67 | 0.34 | 0.50 | 0.40 |
|  | DNNGP | 0.94 | 0.93 | 0.94 | 0.94 |
|  | SoyDNGP | 0.94 | 0.93 | 0.93 | 0.93 |
| PDENS<sup>*</sup> | rDeepGS | 0.85 | 0.84 | 0.83 | 0.83 |
|  | mDeepGS | 0.61 | 0.30 | 0.50 | 0.38 |
|  | DNNGP | 0.84 | 0.83 | 0.83 | 0.83 |
|  | SoyDNGP | 0.85 | 0.84 | 0.85 | 0.84 |
| POD<sup>*</sup> | rDeepGS | 0.82 | 0.77 | 0.69 | 0.71 |
|  | mDeepGS | 0.68 | 0.23 | 0.34 | 0.27 |
|  | DNNGP | 0.80 | 0.76 | 0.66 | 0.69 |
|  | SoyDNGP | 0.83 | 0.80 | 0.70 | 0.73 |
| ST<sup>*</sup> | rDeepGS | 0.81 | 0.71 | 0.63 | 0.64 |
|  | mDeepGS | 0.54 | 0.18 | 0.33 | 0.23 |
|  | DNNGP | 0.80 | 0.67 | 0.61 | 0.61 |
|  | SoyDNGP | 0.82 | 0.68 | 0.63 | 0.63 |

<sup>\*</sup>Flower Color (FC), Pubescence Density (PDENS), Pod Color (POD), and Stem Termination (ST).

### Versatile Predictive Capacity of SoyDNGP Across Diverse Soybean Populations

Our model was developed using the USDA soybean germplasm collections, leaving us uncertain about its application to other resources across diverse countries and latitudes. To appraise the predictive prowess of our constructed models via SoyDNGP, we applied it to a soybean population comprising 559 accessions, inclusive of 121 wild soybeans (*Glycine soja*), 207 landraces, and 231 elite cultivars (Lu *et al*., 2020). We executed predictions for 16 qualitative traits and 12 quantitative traits (Table S2). For substantiating the prediction accuracy for significant yield and quality traits, we juxtaposed the phenotypes of specified soybean traits grown in Zhengzhou, China in 2018 with our predictions. Our analysis unveiled a robust positive correlation between predicted and actual values (Figure 3A). For instance, the correlation for flowering time (R1) and plant height (Hgt) stood at 0.56 and 0.51, respectively. Most impressively, the prediction accuracy for the hundred-seed weight (SdWgt) reached an exceptional 0.84 (Figure 3A). These results suggest that our prediction models bear wide applicability across diverse soybean populations. One potential explanation for the high prediction accuracy for seed weight across different populations could be that environmental factors play a lesser role in this trait compared to others, such as R1 and Hgt (Figure 3A).

**Figure 3.**
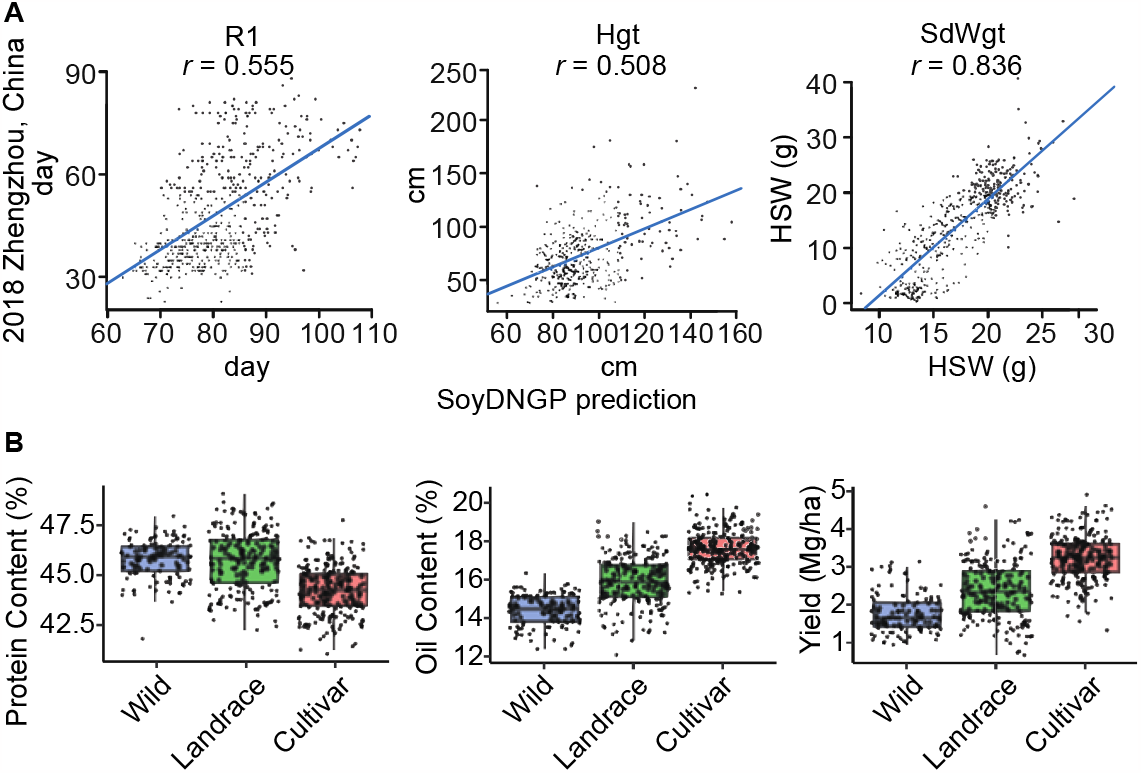
Evaluation of SoyDNGP’s Predictive Capacity in Diverse Soybean Populations. (A) Comparison of observed and predicted phenotypes for selected soybean traits cultivated in Zhengzhou, China in 2018 using the SoyDNGP model. The trend line depicts linear regression. (B) Distribution of predicted phenotypes for a given trait across three distinct subpopulations.

Remarkably, even though wild soybean was not included in model training, our model remains useful for predicting the traits of wild soybean (Figure 3B, Figure S5). For example, our predictions indicated high protein content and lower oil content and yield for wild soybean compared to landrace and elite cultivars (Figure 3B), a finding consistent with prior soybean research (Zhang *et al*., 2018; Swarm *et al*., 2019). This also implies that gene exchange between wild and domesticated soybean might be facilitated by significant gene flow (Wang *et al*., 2019).

### SoyDNGP rooted in soybean but not limit to it

In an effort to evaluate the versatility and efficacy of SoyDNGP, we put it to test with other species, using genotype data and five representative traits from Cotton, Maize, Rice, and Tomato populations (Liu *et al*., 2017; Ma *et al*., 2018; Wang *et al*., 2018; Ye *et al*., 2021). For the sake of comparison, the same datasets were also applied to DNNGP and mDeepGS. Apart from mDeepGS, which exhibited the lowest accuracy, SoyDNGP demonstrated predictive accuracies ranging from an average of 0.50 in Maize to an average of 0.71 in Rice (Figure 4A). A similar performance spectrum was observed in DNNGP (0.49-0.69) (Figure 4A). It’s noteworthy that for smaller sample sizes such as Maize and Tomato, with 214 and 508 samples respectively, DNNGP outperformed SoyDNGP (Figure 4A). However, in larger sample populations like Cotton and Rice, exceeding 1000 samples, SoyDNGP proved superior (Figure 4A). Despite the similarities in accuracy, DNNGP’s mean squared error (mse) was generally higher than that of SoyDNGP (Figure 4B). Based on these findings, we can conclude that SoyDNGP is not only capable of training and predicting phenotypes of traits in other species, but it also surpasses the performance of other methods, thereby confirming its robust versatility and effectiveness. Therefore, SoyDNGP stands as a promising tool for genomic prediction, with its application potentially extending beyond soybeans to other crops and organisms, thereby bolstering the advancements in genomics and breeding research.

**Figure 4.**
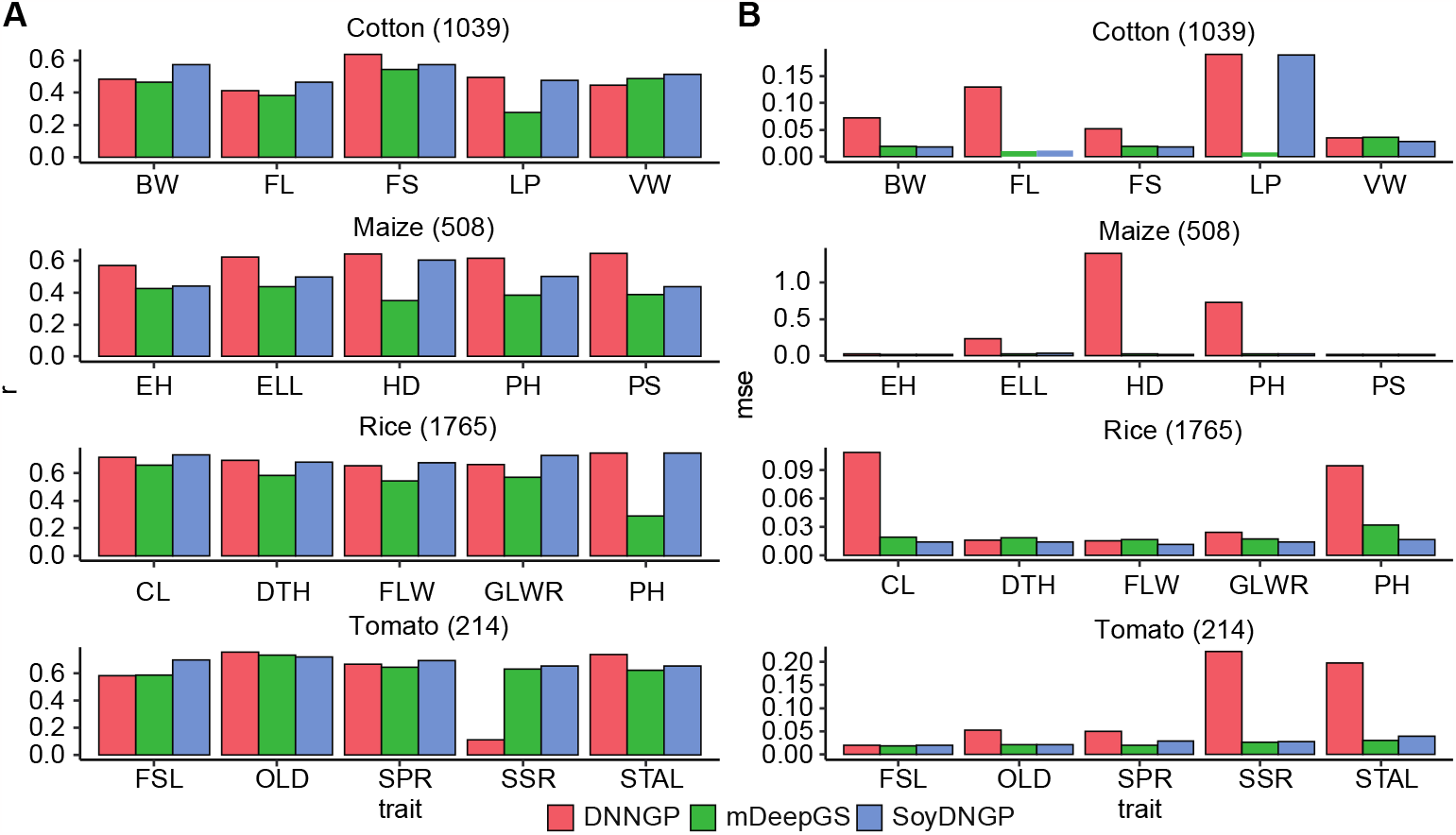
Comparative Evaluation of Predictive Capacity Among SoyDNGP and Two Other Methods Across Diverse Crop Species. (A) Predictive accuracy of various traits assessed by different methods, quantified by the correlation coefficient (r). (B) Absolute errors between normalized observed and predicted phenotypic values for specific traits using different methods. The number in parentheses represents the sample size for a given population. Traits in Cotton include Boll Weight (BW), Fiber Strength (FS), Fiber Length (FL), and Verticillium Wilt (VW). Traits in Maize include Ear Height (EH), Ear Leaf Length (ELL), Heading Date (HD), Plant Height (PH), and Pollen Shed (PS). Traits in Rice include Culm Length (CL), Days to Heading (DTH), Flag Leaf Width (FLW), and Grain Length-Width Ratio (GLWR).

### SoyDNGP is open-friendly web server for the genomic prediction of soybean

To make SoyDNGP accessible to users without deep programming expertise, we have built a web server that bears the same name as our model structure and is available at http://xtlab.hzau.edu.cn/SoyDNGP. The SoyDNGP platform provides two easy-to-use interfaces for exploring trait information. The first feature, ‘Trait Lookup’, lets users enter the taxon identifier, which could be the Plant Introduction (PI) number or traditional name, to check whether the corresponding record is already in our database (Figure 5A). Additionally, our ‘Trait Lookup’ section includes pre-existing trait predictions for 500 soybean accessions, which are in addition to the ones from the USDA soybean germplasm collection, and all have available re-sequencing data. We are continuously increasing this number with daily updates to offer an ever-expanding dataset to our users. This functionality can be beneficial for users wishing to select specific soybean accessions based on certain trait predictions, thereby enhancing the efficiency of SoyDNGP. The second feature, the ‘Trait Prediction’ tool, allows users to upload a Variant Call Format (VCF) file, which our robust predictive models then use to predict trait values(Figure 5A & 5B). We also provide users with the option to contribute to the enrichment of our lookup database. If users opt to contribute, they will not need to rerun the prediction when revisiting their results in the future.

**Figure 5.**
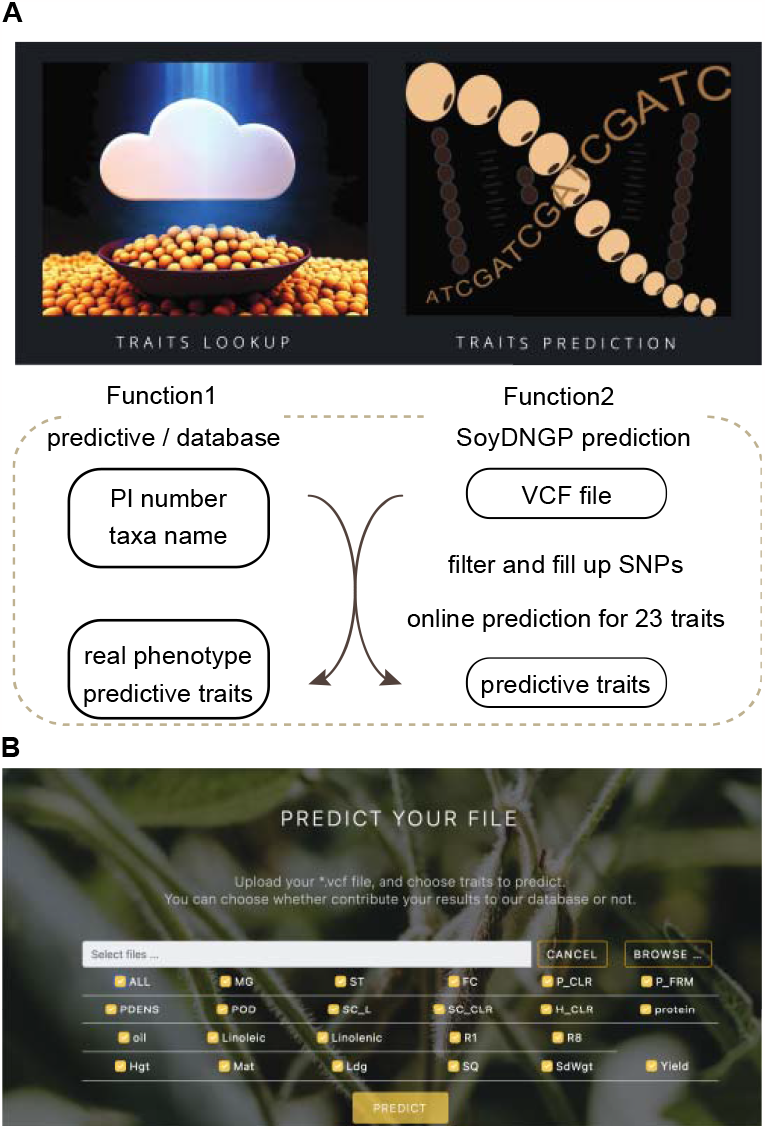
The User Interface and Functionalities of the SoyDNGP Web Server. (A) Descriptions of the two main functions built into the web server are provided. The upper panel showcases the entrance interface on the webpage, while the lower panel outlines the workflow of the two functions. (B) Display of the SoyDNGP prediction function page within the web server. The platform allows users to upload their own VCF files to generate predictions for 28 traits.

## Discussion

SoyDNGP has several significant advantages over DeepGS and DNNGP models:

### Higher Feature Density in Model Input

DeepGS and DNNGP models use a one-dimensional vector, obtained by performing one-hot encoding of the SNP corresponding to the sample, as a feature input for model training (Ma *et al*., 2017; Wang *et al*., 2023). However, this feature has dimension limitations when representing SNP locus feature information in data structure. SoyDNGP uses a three-dimensional matrix as the model’s training input. This structure, while representing genotype information, also considers the relative position information of different SNP loci and the degree of variation difficulty among different types of mutations. As a result, the input feature has higher information density and is more suitable for convolutional neural networks.

### More Rational Model Structure

Both DNNGP and DeepGS use a shallow, wide convolution architecture (Ma *et al*., 2017; Wang *et al*., 2023). While DNNGP adds up to three layers of convolutions on the basis of the DeepGS model, its depth is still insufficient. Shallow wide convolution models use more parameters but have limited deep information extraction and performance abilities, making them less efficient. In contrast, SoyDNGP uses stacking of multiple small convolution kernels, employing a narrow and deep model structure to complete feature extraction. By using as few parameters as possible and introducing more non-linearities, the model ensures more effective feature extraction.

### Reduced Feature Loss

DeepGS employs a 4 × 4 max pooling layer for feature map downsampling (Ma *et al*., 2017). However, due to the high information density of genotype-based feature maps, using a pooling layer would result in considerable feature loss. Hence, SoyDNGP employs a convolution stride of 2 instead of the pooling layer, which achieves feature fusion and downsampling while minimizing feature loss.

### Enhanced Regularization for Stable Model Training

In the DeepGS model, only Dropout layers are used to prevent overfitting and enhance model stability (Ma *et al*., 2017). DNNGP, though it introduces a Batchnorm layer between the first and second convolution layers, still falls short compared to SoyDNGP (Wang *et al*., 2023). SoyDNGP adds Batchnorm and Dropout layers between each convolution layer and incorporates weight decay (L2 regularization) into the optimizer during training, preventing overfitting resulting from the deep network structure (Cortes *et al*., 2012). More Batchnorm layers ensure similar parameter distribution across different convolution layers, making the model training more stable and effective.

### Incorporation of Attention Mechanism

Unlike the CNN models of DNNGP and DeepGS, SoyDNGP introduces a Coordinate Attention mechanism at the beginning and end of the convolution layer. This allows the model to fully consider the feature input’s spatial and channel position information during training, enhancing the relative weight of key points in the feature map and thereby boosting the model’s feature extraction capabilities.

In conclusion, we have created and validated SoyDNGP, a Convolutional Neural Networks (CNN)-based model tailored specifically for predicting soybean traits. We meticulously compared the performance of SoyDNGP with DeepGS and DNNGP models, evaluating predictive accuracy and model complexity. The results underscored that SoyDNGP consistently superseded these models, exhibiting higher accuracy with reduced model complexity. Moreover, we tested SoyDNGP’s adaptability across an array of crop species, including Cotton, Maize, Rice, and Tomato, highlighting its potential as a resilient and versatile tool for genomic prediction. To expand SoyDNGP’s reach, we established a user-friendly web server that offers users easy access to trait predictions and the ability to calculate traits using VCF files. Moving forward, our efforts will concentrate on consistently augmenting the database of pre-existing trait predictions and enhancing the accuracy and efficiency of the model. With ongoing learning and development, SoyDNGP is poised to make significant contributions to the progress of soybean breeding and potentially broader agricultural advancements.

## Acknowledgments

We express our gratitude to Dr. Ting Zhao of Zhejiang University for the provision of cotton population genotype and phenotype data; we are also thankful to Dr. Xin Wang and Dr. Ning Yang from Huazhong Agricultural University for their contribution of tomato and maize data sets. This research was made possible through the support of the National Key Research and Development Program of China (grant 2022YFD1201502).

## Author contributions

The research was conceptualized by XW, while the population data and key trait information were supplied by CF and FK. The data analysis was jointly undertaken by XW, PG, HZ, ZL, YFL, and YLL. The development of the SoyDNGP model structure and web server were spearheaded by PG and HZ, respectively. The manuscript was primarily penned by XW.

## Data availability

Complete data sets can be found within the main text, supplementary materials, and referenced studies, as well as in public databases. The code, pre-built models, and the standalone network structure are accessible at https://github.com/IndigoFloyd/SoybeanWebsite

