## Supplementary material for "SoyDNGP: A Web-Accessible Deep Learning Framework for Genomic Prediction in Soybean Breeding": Figure S1-S5, Table S1

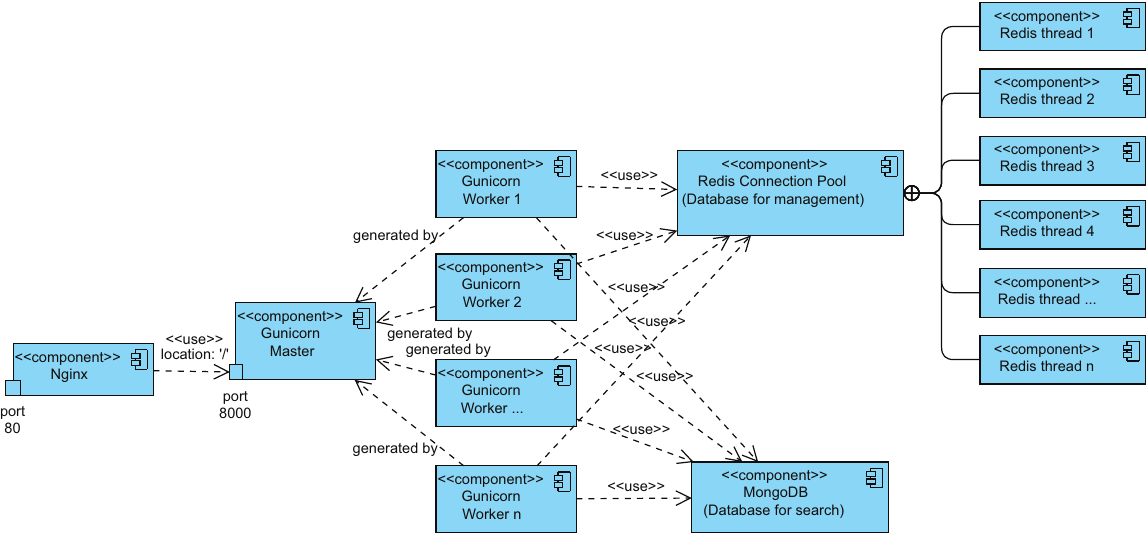

**Figure S1. The Components of SoyDNGP’s Web Server.**

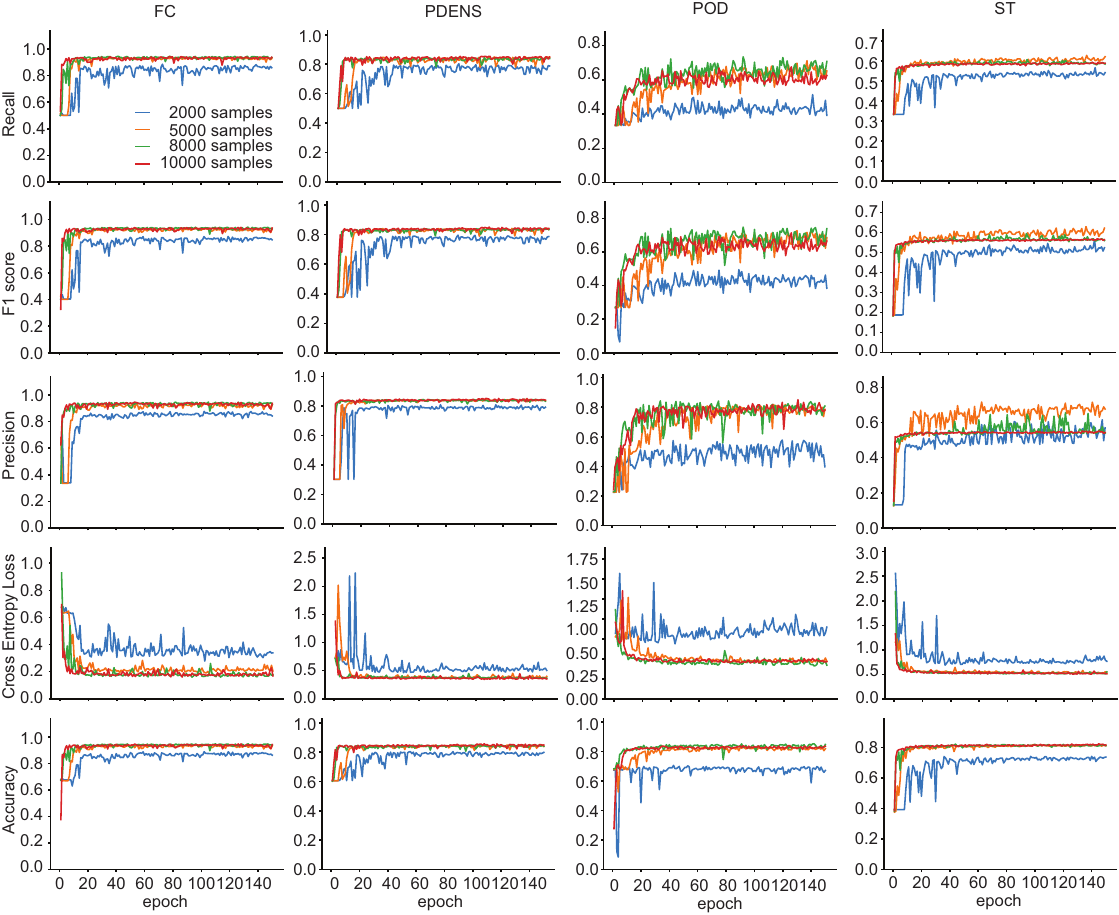

**Figure S2. Assessment of SoyDNGP's predictive performance in classification tasks with varying sample sizes.** FC, PDENS, POD, and ST signify Flower color, Pubescence density, Pod color, and Stem termination, respectively.

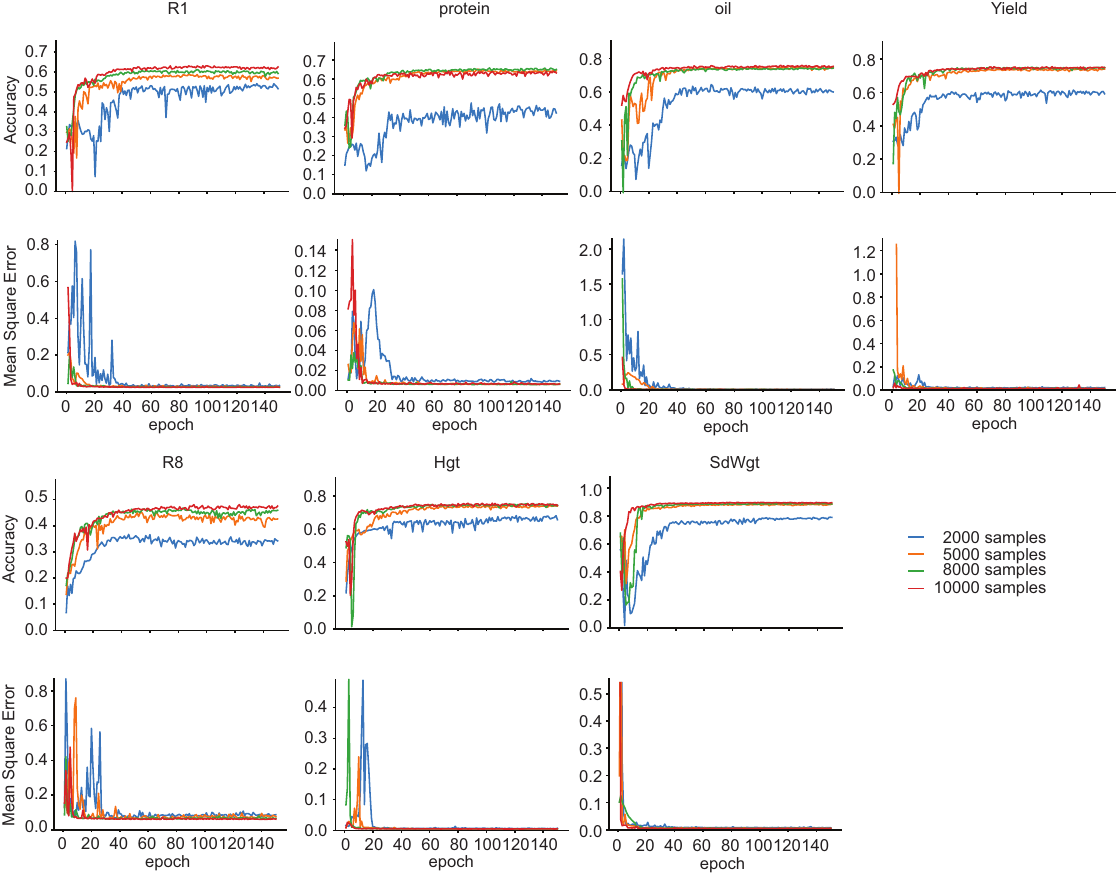

**Figure S3. Assessment of SoyDNGP's predictive performance in classification tasks with varying sample sizes.** Accuracy is quantified by the correlation coefficient (r). R1 represents Flowering Time.

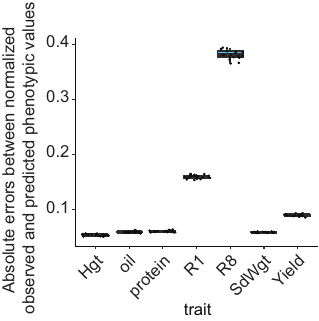

Top of Form

**Bottom of Form**

**Figure S4. The absolute errors between the normalized observed and predicted phenotypic values.** Hgt, Oil, Protein, R1, R8, SdWgt, and Yield stand for Plant height, Oil content, Protein content, Flowering time, Maturity time Hundred seed weight, and yield, respectively.

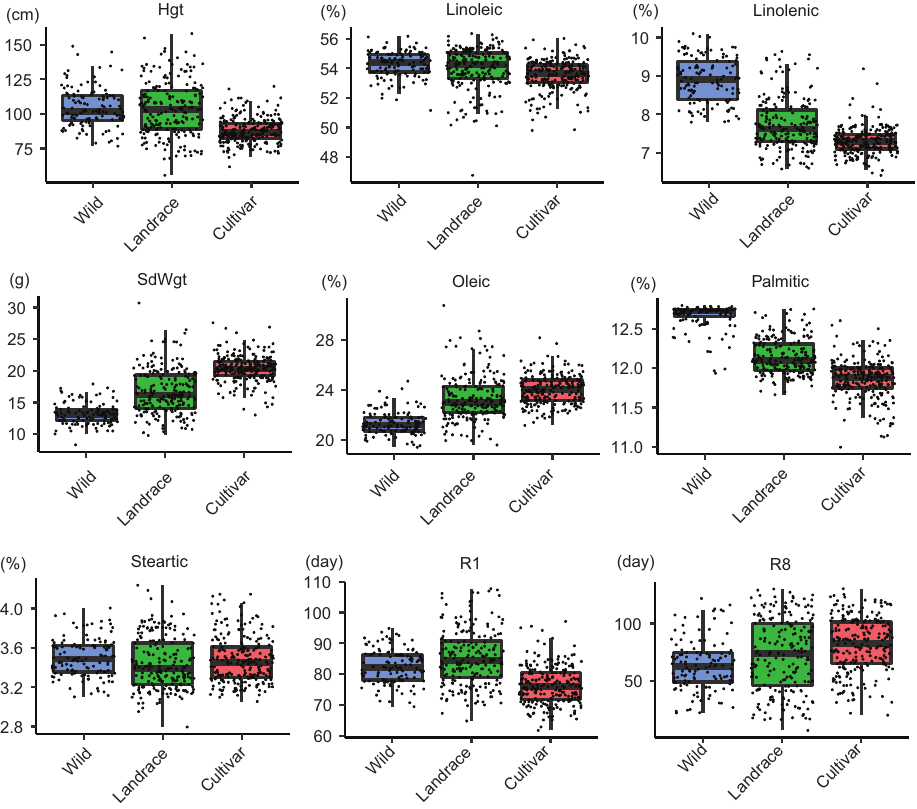

Figure S5. Distribution of predicted phenotypes for a given trait across three distinct subpopulations.

Table S1. the total parameter count in the four methods.

| **methods** | **total parameter mount** |
| --- | --- |
| rDeepGS | 6,149,048 |
| mDeepGS | 9,456,352 |
| DNNGP | 3,076,736 |
| SoyDNGP | 69,536 |
